# Optimization and validation of a Protein Phosphatase inhibition assay for accessible microcystin detection

**DOI:** 10.1101/2022.08.16.502937

**Authors:** JE Alba Posse, C Gonzalez, P Carriquiriborde, A Nadra, J Gasulla

## Abstract

The presence of cyanobacterial toxins in freshwater constitutes an increasing public health concern, especially affecting developing countries where the high cost of available methods makes monitoring programs difficult. The phosphatase inhibition assay (PPIAs) is a sensitive method with low instrument requirements that allows the quantification of the most frequent cyanotoxins, microcystins (MC). In this work, we implemented a PPIAs, starting from Protein Phosphatase 1 (PP1) expression up to the validation with samples of algal blooms from Argentina. To do this, we optimized the expression and lyophilization of PP1, and the assay conditions. Also, we included robustness and possible interfering analysis. We evaluated the most widely used cyanobacterial lysis methods and determined that heating for 15 minutes at 95°C is simple and adequate for this assay. Then, we performed MC spikes recovery assays on water samples from three dams from Argentina, resulting in a recovery ranging from 77 to 115%. The limit of detection (LOD) was 0.4 μg/L and the linear range is 0.4 μg/L - 5 μg/L. Finally, we evaluated 64 environmental samples where MC was measured by ELISA test containing from 0 μg/L to 625 μg/L. The PPIA showed excellent correlation (Pearson correlation coefficient = 0.967), no false negative and no false positives above the 1 μg/L WHO guideline (0.11 false positive rate). In conclusion, we optimized and validated a PPIAs to be an effective and accessible alternative to available commercial tests.

## Introduction

### 1.1. A growing concern: microcystin cyanotoxin

Cyanobacteria are microscopic photoautotrophic prokaryotes that inhabit most water bodies. They are the base of the trophic pyramid and are responsible for many essential functions in the biosphere, such as nitrogen fixation and oxygen production. However, under certain conditions highly related to eutrophication and anthropic footprint (such as domestic and industrial effluents discharges, misuse of agrochemicals, and global warming), many of them can produce a wide variety of toxic metabolites, the cyanotoxins. Therefore, the incidence of cyanobacterial harmful algal blooms (*CyanoHABs*) in freshwater bodies has become a worldwide issue of increasing concern (Berman, 2022; O’Farrell et al., 2019; O’Neil et al., 2012).

Cyanobacterial hepatotoxins Microcystins (MC), are cyclic non-ribosomal heptapeptides of ~1 kDa, with more than 250 reported variants (Foss et al., 2018). They are the most ubiquitous group of toxic cyanobacterial metabolites, affecting almost every region of the globe. Hence, they have been studied for years due to their carcinogenic and hepatotoxic effects (Rastogi et al., 2014). One of the most critical incidents in the last years involved the use of untreated water affected by an algal bloom, in a hemodialysis clinic in Caruaru, Brazil. It resulted in 101 cases of acute liver injury and 50 deaths due to liver failure (Jochimsen et al., 1998). Several other reports have been made describing their toxicity for humans and animals, in many degrees of chronic and acute exposure (Andrinolo et al., 2008; Foss et al., 2018; Giannuzzi et al., 2011; Melaram et al., 2022; Mez et al., 1997; Tamele and Vasconcelos, 2020; Wang et al., 2010). Furthermore, ecosystems can also be affected by MC. Several episodes of MC bioaccumulation and biomagnification in animal tissue were reported (Miller et al., 2010; Pham and Utsumi, 2018). Therefore, their quick and precise detection is relevant to public health and environmental preservation. Additionally, specialists suggest that the current lack of active monitoring of cyanotoxins could be leading to the underestimation of the actual number of water poisoning episodes related to MC (Aguilera et al., 2018; Tamele and Vasconcelos, 2020). This is an especially concerning aspect for developing countries, where resources to make this monitoring possible are mostly limited. In some cases, this restriction occurs due to the elevated cost of the detection equipment or detection kits (Aguilera et al., 2018; Falconer, 2001; Meneely et al., 2018), which are generally imported from developed countries.

### 1.2. Detection of Microcystins

The first step of standard detection protocols is the cyanobacterial lysis, to get the full release of toxins. This is usually achieved by successive cycles of freeze-thawing of the sample. There are also alternative MC purification protocols, where cyanobacteria are arduously separated from samples by nylon filtering followed by organic extraction of metabolites (Sevilla et al., 2009). The “Gold standard” for MC determination is through HPLC (High-performance liquid chromatography), with a previous clean up step that includes filtration with a series of glass fiber and membrane filters, followed by a concentration step in methanol with solid phase extraction C18 ODS cartridges (Harada et al., 1988; Lawton et al., 1994). Finally, detection is performed by HPLC-MALDI-TOF (Matrix-Assisted Laser Desorption/Ionization, Time of Flight), LC-MS (Liquid chromatography-mass spectrometry), among other techniques (Kumar et al., 2020; Singh et al., 2012). Detection limits are approximately 0.1 μg MC/L. However, the implementation of these high-budget detection methods in water samples is still difficult for developing countries due to their high-cost equipment and specialized personal requirements (Catanante et al., 2015; Tamele and Vasconcelos, 2020).

Only a few alternative approaches can stand up to the standards established by the WHO (World Health Organization) for chronic exposure in freshwater of 1 μg MC/L (Graham, 1999), guideline values discussed in (Hitzfeld et al., 2000). Among those, *Enzyme-Linked ImmunoSorbent Assay (ELISA*) methods are some of the most popular, and actually have commercial implementations that allow *in-situ* measurements (*Abraxis Strip Test kit™*) (Humpage et al., 2012). ELISA-based systems usually present cross-reactivity between toxin variants, sometimes detecting high concentrations of non-toxic variants, but underestimating the most toxic ones (Alvarenga et al., 2014; Carmichael and An, 1999; Moore et al., 2016). Also, as mentioned in (Humpage et al., 2012), they tend to over-respond to the concentrations in the lower range segment (2,5 μg/L-5 μg/L). Also, MC can be detected indirectly by measuring the presence of genes of the *myc* cluster in the water sample, instead of the toxin itself (Pacheco et al., 2016). Another widely known approach is the PPIAS (*Protein Phosphatase Inhibition Assays*) (An and Carmichael, 1994).

### 1.3. PPIAS

PPIAs are detection tests substantiated in the selective inhibition of the phosphatase activity of Ser-Thr-phosphatase proteins by MC. Usually, they couple protein activity with a chromogenic substrate, but electrochemical detectors were also developed (Campàs et al., 2008; Catanante et al., 2015). Although in most cases the product is the yellow pNP (absorbing at 405 nm), some papers reported the usage of Malachite Green as an activity reporter (Heresztyn and Nicholson, 2001). In the last years, several efforts have been done to improve PPIAs (Heresztyn and Nicholson, 2001; Sassolas et al., 2011; Garibo et al., 2014; Rapala et al., 2002; Ward et al., 1997; Zhang et al., 1992). PPIAS-based commercial test kits are available in the market as Microcystest™ and ZeuLab.

### 1.4. Protein Phosphatase 1

Protein Phosphatase 1 (PP1) is a metal-regulated monomeric and globular 38 kDa serine/threonine phosphatase from the phosphoprotein phosphatase (PPP) family, which is ubiquitous among eukaryotes (Barford, 1996; Goldberg et al., 1995; Heroes et al., 2015, 2013; Shi, 2009). It is widely known that the phosphatase activity of PP1 is selectively inhibited by the interaction with certain types of cyanobacterial hepatotoxins, being MC (and particularly MC-LR) the most prevalent and toxic. Previously PP1 was purified from rabbit muscle homogenate (Cohen et al., 1988). Nowadays PP1 can be expressed in *Escherichia coli* and purified by a tag (Sassolas et al., 2011). However, the recombinant expression and purification of PP1 in bacteria leads to low expression yields, mainly to misfolding and relocalization to inclusion bodies (Zhang et al., 1992). Some authors, interested in PP1 structural characterization, reported that coexpression with GroEl/Es chaperones with an in-vivo refolding step improved substantially the concentration of soluble PP1 obtained in *E. coli* DE3 (BL21) (Kelker et al., 2009; Peti et al., 2013).

### 1.5. Objectives

Considering the international need for accessing cheaper MC detection methodologies that allow systematic and expanded monitoring in the territory, we intended to optimize the conditions for the development of a low-cost microcystin PPIA biosensor, based on the selective inhibition of PP1. In this work, we approach the problem covering the entire process. From the optimization of PP1 expression and storage conditions, to the validation with environmental samples of algal blooms and *Microcystis aeruginosa* cultures, including studying the robustness of the test conditions, assay’s interferences, and evaluating lysis protocols.

## Methodology

### 2.1. Salts and reagents

Solid p-Nitrophenyl Phosphate (pNpp) was purchased from Santa Cruz Biotechnology (California, USA). Bovine Serum Albumin (BSA) was purchased from Sigma Aldrich (Misuri, USA). NTA-Ni Histidine trap beads were purchased from GE Healthcare (Illinois, USA). The remaining metal salts, buffer and medium components were obtained from Biopack and Cicarelli (Argentina).

### 2.1. Protein expression and purification

We double-transformed *Escherichia coli* BL21(DE3). We used a RP1B-PP1ca (pET28 family plasmid, AddGene catalog number #51768) with kanamycin resistance, expressing the rabbit PP1α catalytic subunit (aa 7-300) linked to a histidine tag for purification. The second vector, kindly provided by Dr Diana Wetzler, presented chloramphenicol resistance and contained the expressing sequence for both chaperones GroEL and GroES. The sequences of the chaperones and PP1 were confirmed by Sanger sequencing. Both PP1 and chaperones were under T7 IPTG/lactose-induced promoters. Transformed bacteria were grown overnight in solid Luria Bertani (LB) medium plaques with both antibiotics. Plasmid incorporation was corroborated by MiniPrep and agarose gel electrophoresis. For protein expression, we proceeded according to an adapted version of previous protocols (Kelker et al., 2009; Peti et al., 2013). First 10 mL pre cultures of LB medium with both antibiotics and 1mM MnCl_2_ were incubated overnight and added the next day to a 1L LB medium flask with antibiotics (25 μg/mL chloramphenicol, 50 μg/mL kanamycin) and 1mM MnCl_2_. The culture was grown at 37 °C under agitation (200 RPM) until OD = 0.6 was reached. Then culture expression was induced with 10 g of lactose and the temperature was changed to 28°C in agitation for 6 hours. After that, cells were harvested by centrifugation and resuspended in 50 ml of LB medium with 250 μg/mL chloramphenicol, 1mM MnCl_2_, 50 μL protease inhibitor PMSF, and 50 μg/mL kanamycin. Cells were incubated in agitation at 200 RPM at 15 °C for 2 hours in an *in-vivo* refolding step as a modified version of (Peti et al., 2013). Cells were harvested again by centrifugation in 50 mL Eppendorf tubes and pellets were stored at −20°C. The next day cells were resuspended in 25 mL of buffer (pH 8, 700 mM NaCl, 1mM MnCl_2_, 50mM Tris, 1 mM DTT, 0.1% Triton X100) and lysed in ice by sonication (Sonics Vibra-cell, Connecticut, USA) in 8 cycles of 30/60 seconds (ON/OFF) cycles at 30 kHz. The lysate was centrifuged two times for 20 minutes, at 4°C, at top speed velocity (17000 g) (Beckman JA-20 Rotor, Indiana, USA), and the supernatant was conserved.

Protein was purified in the same buffer with lower ionic strength (70 mM NaCl instead), through an affinity Nickel-NTA Agarose Resin (Gold Biotechnology, Missouri, United States) column by gravity. After loading the protein into the column, we performed a washing step with 50 column volumes of 20mM imidazole. After that, the column was eluted slowly with incrementing imidazole concentrations ranging between 50 mM to 1M. Fractions were analyzed in a 14% SDS PAGE revealed by Coomassie Blue. PP1 appeared without contaminations in fractions of 250 mM of imidazole. Concentrations in every fraction were measured by Bradford method and read in a multiwell plate reader (TECAN INFINITE PRO, Switzerland) at 595 nm. For preservation, the protein was gently mixed with glycerol (20% final concentration), fractionated in 500 μL Eppendorf tubes and instantly frozen in liquid nitrogen and stored in a −80°C Freezer (UltraFreezer Righi, Argentina).

### 2.2. Phosphatase Activity Determination

The reaction buffer consisted of 40mM Tris HCl, 20 mM KCl, 30 mM MgCl_2_, and 2 mM MnCl_2_, pH = 8, as reported by (Carmichael and An, 1999). Protein activity was measured in a colorimetric assay on 96 well plates at a final volume of 200 μL. First 50 μL of protein diluted in *reaction buffer*, were mixed with 100 μL of *reaction buffer*, and 50 μL of substrate solution (*reaction buffer* supplemented with 12 mg/mL of p-nitrophenyl phosphate (pNPP), 3 mM DTT) were added to start the reaction. The plate was incubated at 37°C while absorbance was read at 405 nm every 5 minutes for 1 hour in a multi-well plate reader (*TECAN INFINITE PRO, Switzerland*). The background activity measured from a blank solution without protein was always subtracted from the reported activity.

#### 2.2.1. Protein activity inhibition in solution

A MC-LR solution (Microcystin LR Standard 0.5 mg Eurofins Abraxis catalog number #300630, stored in 100 mg/L methanol stock solution) was diluted in calibration curves from 0.01 μg/L to 3000 μg/L (final concentration of methanol 8%). The effect of methanol in PP1 was tested and no significant inhibitory effect was found up to 8% (Supplementary Figure 1C). For the phosphatase inhibition assays, 50 μL of diluted PP1 were incubated 10 minutes with 100 μL of MC-LR standard of each concentration of the curve (instead of buffer), by triplicate. Coefficient of variation % (CV%) were calculated to ensure they do not exceed 20% as indicated by (Carmichael and An, 1999). For pH assays, protein solutions were prepared with buffers previously adjusted to pH 7, 8, 8.6, and 9 respectively. For salinity activity tests, protein solutions were prepared in a buffer with 0 mM, 50 mM, 100 mM, and 200 mM of NaCl, from a 4M NaCl stock. For interference inhibition assays we measured the activity of PP1 in the presence of 16 different salts, in concentration curves ranging from 10 μg/L to 300 mg/L (of the solid salt of the following compounds: *Ag_2_SO_4_, Cu_2_SO_4_, CaCl_2_, CuCl_2_, MgCl_2_, MgSO_4_, MnCl_2_, MnSO_4_, Na_2_CO_3_, NaF, Na_2_AsO_4_, Na_3_PO_4_, NiSO_4_, Na_3_VO_4_, PbNO_3_, PbCl_2_*). Curves were performed in 40 mM Tris buffer pH=8, unless cation solubility was severely compromised by precipitation at buffer pH (*Ag_2_SO_4_, Cu_2_SO_4_, CuCl_2_*). In those cases, curves were performed in bidistilled water.

#### 2.3.1 Protein storing conditions

The protein was purified and quantified the same day and was aliquoted in purification buffer, in 0.5 mL Eppendorf tubes with 300 μl PP1 solution with the addition of 20% glycerol and stored in different conditions: 4°C, −20°C, −80°C. For each measurement, three tubes were taken out from each storage treatment every week. All tubes were centrifuged and both activity and mass (Bradford method, with BSA 0-200 μg/mL calibration curve) were determined. For lyophilization assay, fresh protein was diluted 1/200 in *reaction buffer* with 2 mM MnCl_2_, and 50 μl of protein were added to each well in a 96 multiwell plate. As preservants, 50 μl of *reaction buffer*, 28 μl of 50% trehalose and 12 μL of BSA 20 mg/mL were added to each well. For the blank, 50 μl of *reaction buffer* was added instead of the protein. The lyophilization occurred at 4 Pa O.N. in a freeze dryer (Biobase, China). Lyophilizate was carefully covered with an aluminum adhesive tape and kept at −20 °C. Each week protein was rehydrated with 150 μl of bidistilled water and 50 μl of reaction buffer with the substrate was added to determine activity. The activity was relativized to the initial value.

#### 2.3.1 Microcystis isolation and culture

Fresh phytoplankton material was collected from Rio de la Plata, Argentina, during a cyanobacterial bloom. Samples potentially containing microcystins were used for the isolation of *Microcystis aeruginosa* (Kützing) organisms. The isolation of individual colonies from the sample was carried out using the glass capillary method described by (Andersen and Kawachi, 2005) under an optical microscope. The isolated organisms were cultured in glass bottles containing 150 mL of BG11 medium (Rippka et al., 1979) at 24°C with a 16:8 (light:dark) photoperiod under fluorescent light. *Microcystis* colonies were harvested from the monocultures in the exponential phase (Andersen and Kawachi, 2005) for running the experiments.

#### 2.3.2 Cyanobacterial lysis

Starting from 2 mL samples of culture, we tested 3 lysis treatments and the control without treatment. After each treatment, samples were centrifuged at 13500 g for 10 minutes, and the supernatant was kept. As an index of cellular rupture, we measured the *Internal Organic Matter Release* (IOMR) (Geada et al., 2019) by UV absorbance at 254 nm in a UV-VIS spectrophotometer (Jasco™, Japan). Toxin release into the solution was measured by PP1 inhibition assay. The results shown are the average of 6 independent assays, each one evaluated in 3 technical replicates. Additionally, two samples of the culture were treated according to (Sevilla et al., 2009) with minor modifications. Briefly, 20 mL of water was filtered through a nylon 0.45 μm membrane. The filtrate was used to determine MC-LR (indicated as extracellular MC, ECMC). The filter was incubated 15 minutes in 5 mL of 100% methanol twice. The two extractions were pooled and diluted with the necessary distilled water for toxin measurement. MC-LR was measured in the extract by PP1 inhibition assay (indicated as intracellular MC, ICMC). The total MC (TMC) was calculated as the sum of both ECMC and ICMC. For the *heating* treatment, we incubated the samples for 15 minutes in a 95°C heating block (*AccuBlock, Labnet Inc., Virginia, USA*). Some authors indicated that freeze-thawing at −20°C in 2 hours cycles results in inefficient cyanobacterial lysis (Pestana et al., 2014). Hence, *freeze-thaw cycle* treatment was performed in 3 cycles of overnight freezing, where aliquots were stored at −20°C for more than 12 hours and then thawed, as proceeded by previous authors and standard ELISA protocols (Geada et al., 2019). Sonication was done on an ice bath, for 2 minutes in cycles of 10/30 seconds (ON/OFF) at 30 kHz.

### 2.4. Determination of microcystin-LR using LC-ESI-MS/MS

The *Microcystis* culture sample was held for 72hs at −20°C to release the intracellular toxin from cyanobacteria cells. After thawing, the culture was passed through an Polyethersulfone (PES) 0.45 μm syringe filter to extract the MC-LR toxin. Because of the high toxin concentration detected previously, a 1:1000 dilution was prepared and collected in a 2 mL vial. A calibration curve was carried out using five levels with Abraxis™ MC-LR Standard solution (R-Square: 0.9977,Supplementary Figure 3A). Liquid chromatography was performed based on EPA Method 544. Chromatographic separation was performed using the Thermo Scientific™ Vanquish™ Core HPLC system equipped with a Thermo Scientific™ Accucore™ C18 LC column (2.6 × 100 mm, 2.6 μm) maintained at 30°C. The mobile phase was Water (0.05 formic acid + 0.05 TFA)-Acetonitrile. The injection volume was 5 μL.

The MC-LR was separated and identified by comparing the acquired mass spectrum and retention time to the reference spectrum and retention time for calibration standard acquired under the same LC-MS/MS conditions (Haghani et al., 2017). MC-LR was detected on a Thermo Scientific™ TSQ Fortis™ triple quadrupole mass spectrometer equipped with a Heated Electrospray Ionization source (HESI). MC-LR concentration was determined by calibration curve comparison. The MC-LR final concentration was obtained as chromatogram MC-LR amount (μg/L) * Fdil=1000.

### 2.5. Measurement of field water samples and spike detection

Freshwater samples were taken and immediately sent refrigerated to us by our collaborators: Sampling was done within monitoring programs by the local authorities of the corresponding basins: Salto Grande dam, Entre Ríos province, Argentina; El Chocon dam, Neuquén province, Argentina; Río De La Plata, Buenos Aires province, Argentina (Supplementary Table 1). One of the MC-free samples (Supplementary Table 1, Sample ID# 59) was divided into six 2 mL Eppendorf tubes (n=6). Each tube was spiked with 0, 1, 3, or 10 μg/L of MC-LR, final concentrations. The tubes were treated for 15 minutes in a 95°C heating block and then centrifuged at maximum speed. MC concentration was measured by our PP1 inhibition assay, and Recovery % was calculated. For the matrix-effect assay, we performed the same protocol with 3 replicates for the other 7 samples where no MC was detected (Supplementary Table 1 Samples ID# 1-7).

### 2.6. Statistical analysis

For inhibition curves data was fitted to a sigmoidal 4-parameter curve, and inhibition concentration 50 (IC50), Top, Bottom, and Hill-coefficient were estimated in *GraphPad Prism 8.0*. For comparative analysis between treatments, t-student tests and ANOVA tests were performed. Homoscedasticity (Levene Test) and residue normality (Shapiro Wilks Tests) were tested with *RStudio* when applicable. The significance criterion was considered *p*<0.05. Error bars represent standard error of the mean (SEM) unless otherwise stated.

## Results

### 3.1. Protein expression, purification, and storage

The protein was successfully expressed and purified. Its interaction with the highly concentrated chaperones was considerable and, at first, resulted in a coelution between PP1 and chaperones. Reducing ionic strength of the elution buffer from (700 mM to 70 mM NaCl) combined with performing a large-volume cleaning step (with 20 mM Imidazole), and a slow column elution, improved the purity of the PP1 obtained (Figure 1A).

**Figure 1.**
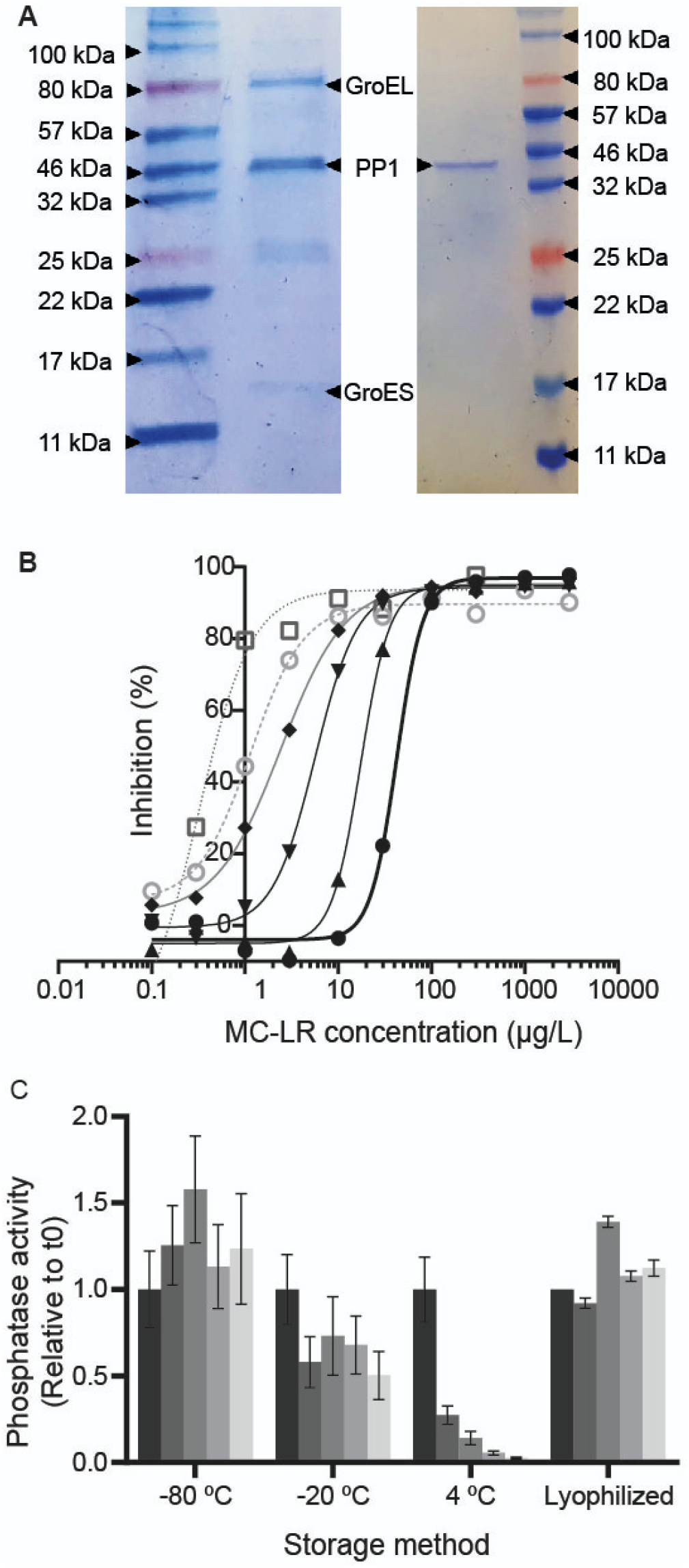
Expression, characterization and storage of PP1. A. SDS-PAGE showing protein purification after eluting from NTA-His column, against molecular marker (*NEB 1 Kb*) Left: Chaperones coeluted with PP1. Right: PP1 was purified from chaperones due to extensive washing with 20mM Imidazole. B. Inhibition profile displacement while diluting protein from 62.5 μg/mL (filled circles and thick line) in six exponential half dilutions down to 1.95 μg/mL (empty square and dotted line). C. Phosphatase activity decayes in time for four storage conditions. The left dark-gray bar of each treatment corresponds to t=0. The following gray bars correspond to weeks 1-4. Error bars correspond to SEM.

The inhibitory response was tested with MC curves (0.1 - 3000 μg/L MC-LR). As expected, the observed sensitivity was highly dependent on the protein concentration. A series of 6 dilutions (1:2) were made to a protein of 62.5 μg PP1 /mL down to 1.95 μg PP1/mL. As seen in Figure 1B, the inhibition curve displaces to lower IC50 as protein concentration diminishes, enhancing sensitivity but compromising signal intensity. The inhibitory concentration 50 (IC50) of the corresponding dilutions were (μg/L MC-LR): 42.3; 17.37; 5.65; 2.42; 1.15; and 0.31. For each of the following experiments, protein concentration was adjusted between 2-10 μg PP1/mL, to maintain the IC50 in a range between 1-3 μg MC/L. The protein was aliquoted on the same day of purification and different storage conditions were evaluated (4°C, −20°C, −80°C). The control treatment activity decayed quickly at 4°C (73% of decay after the first week) and resulted in no measurable activity remaining after 1 month of storage. At −20°C, the measured activity diminished to a mean of about half the initial value after 4 weeks. Nevertheless, the reduction was not statistically significant significantly (*p*=0.06, ANOVA). The concentration of protein at −20°C also diminished in a comparable proportion (*data not shown*). For −80°C, no significant decay in activity or mass was seen after 4 weeks (*p*>0.05, ANOVA) (Figure 1C). Moreover, the PP1 was lyophilized after purification directly on 96 wells plates. The protein activity was measured each week after the freeze-drying for one month. As seen in Figure 1C, protein phosphatase activity remained unaltered in the chosen period. The same process was tested to evaluate if the MCLR curve could be lyophilized in the same well as the protein. As seen in Supplementary Figure 1B, lyophilized MCLR exhibited less inhibition than fresh MCLR, suggesting that the lyophilization process is degrading some amount of toxin.

### 3.2. Effect of salinity and pH on PP1 activity and MC sensitivity

Freshwater samples present a certain degree of variability referring to salinity. In order to ensure assay robustness both assay sensitivity and signal intensity (protein activity) must be tolerant to water with all ranges of salinity for freshwater (up to 0.5 g NaCl/L, or 8 mM NaCl) (“Salinity of Water,” n.d.), and some range of brackish (up to 10 g NaCl/L). The protein presented a stable response at the different concentrations of salt evaluated in this work (up to 200 nM NaCl). Neither the sensitivity nor activity (Figure 2A, 2B respectively) varied significantly among the NaCl gradient compared to the control treatment without NaCl (*ANOVA, p>0.05*). In seawater, salinity exceeds 35 g NaCl/L (600 mM NaCl) (“Salinity of Water,” n.d.). Activity assays performed at higher salinities showed a diminishing of 50% of activity (*p*<0.05) when tested in NaCl higher than 1M (Supplementary Figure 1A). To ensure valid results in those conditions further studies are needed.

**Figure 2.**
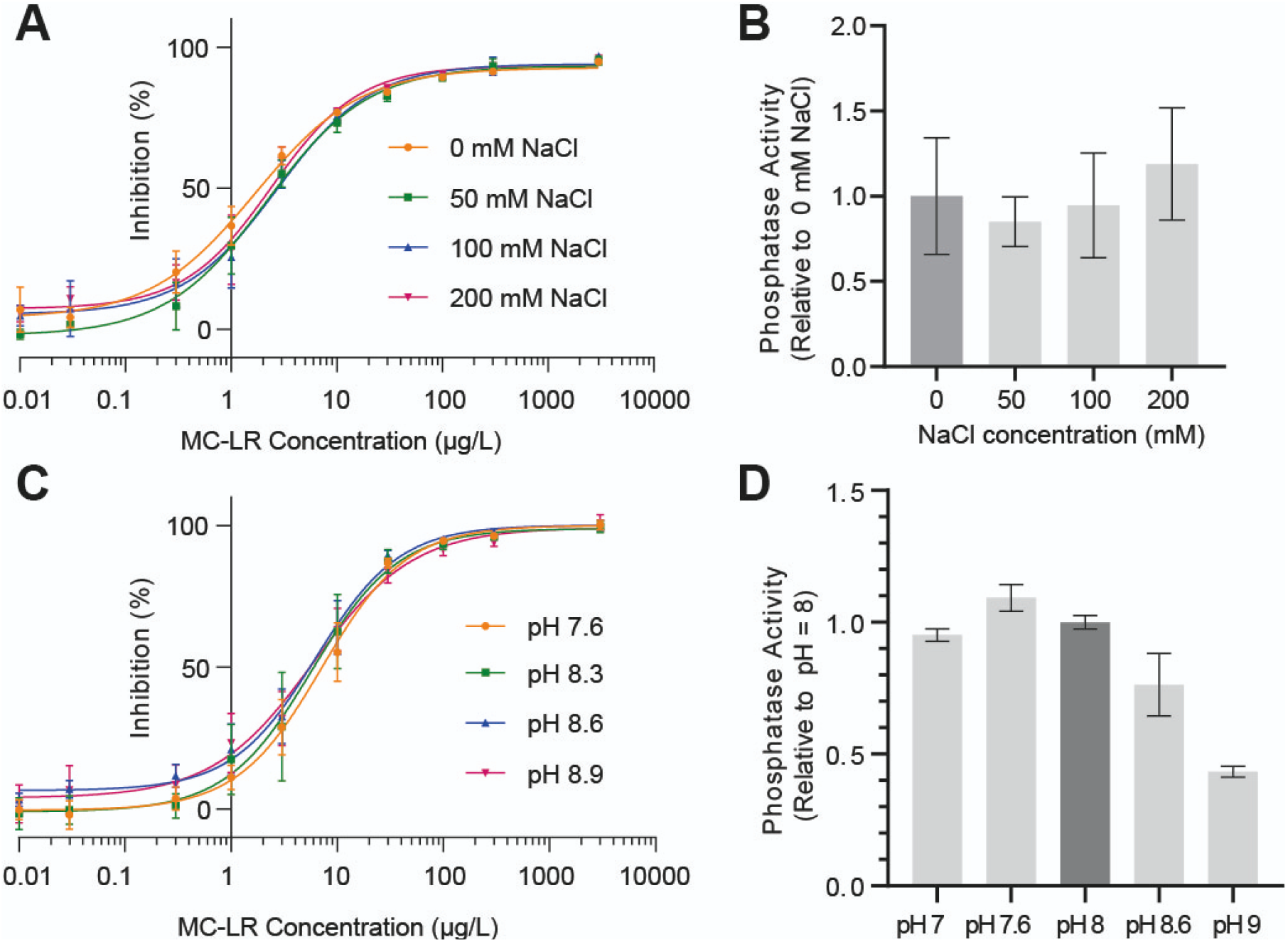
Robustness of the PP1 inhibition assay. A. Phosphatase activity Inhibition in a gradient of MC-LR for different NaCl concentrations. B. Phosphatase activity reported at different NaCl concentrations relativized to 0 mM NaCl. C. Phosphatase activity Inhibition in a gradient of MC-LR for different reaction pH. D. Phosphatase activity reported at different reaction pH relativized to pH 8.

Environmental samples report a wide spectrum of pH, but the main conditions in which *Microcystis aeruginosa* grows and thrives among other genera is between pH 7-9 (Yang et al., 2018). Also, *Microcystis* is incapable of growing neither in pH below 4 nor above 12. (Wang et al., 2011). Furthermore, the PP1 has Mn^2+^ ions as cofactor, whose solubility is severely diminished in pH superior to 8 due to oxidation with the air (Ghosh and Agate, 2015), compromising enzymatic activity. Hence, it was relevant to quantify the effect of pH in both sensitivity and signal intensity in the range of pH 7-9. Protein activity was measured at different pHs (Figure 2D, activity relativized to pH = 8). As observed, the activity significantly decreased for pHs higher than 8. However, IC50 at pH inhibition curves (Figure 2C) presented no significant alterations for the mentioned pHs (IC50 with *p*>0.05). The maximum activity is presented at pH 7,6. This is consistent with previous works (Heresztyn and Nicholson, 2001).

### 3.3. Chemical interference

PP1 is a metal-coordinated protein and thus, other cations can interfere with it, varying the resultant activity. To avoid toxin misestimation we quantified the inhibitory effect of many cations and anions salts, including the most prevalent salts in water bodies, and the ones we considered relevant according to bibliography. To achieve this, we constructed PP1 inhibition curves for different combinations of ions. In Figure 3, we present only those that produce results relevant to be discussed: NiSO_4_, Na_3_VO_4_, Na_3_PO_4_, Cu_2_SO_4_ and Ag_2_SO_4_. The rest of the curves, with an effect of ≤ 25 % are presented in Supplementary Figure 2.

**Figure 3.**
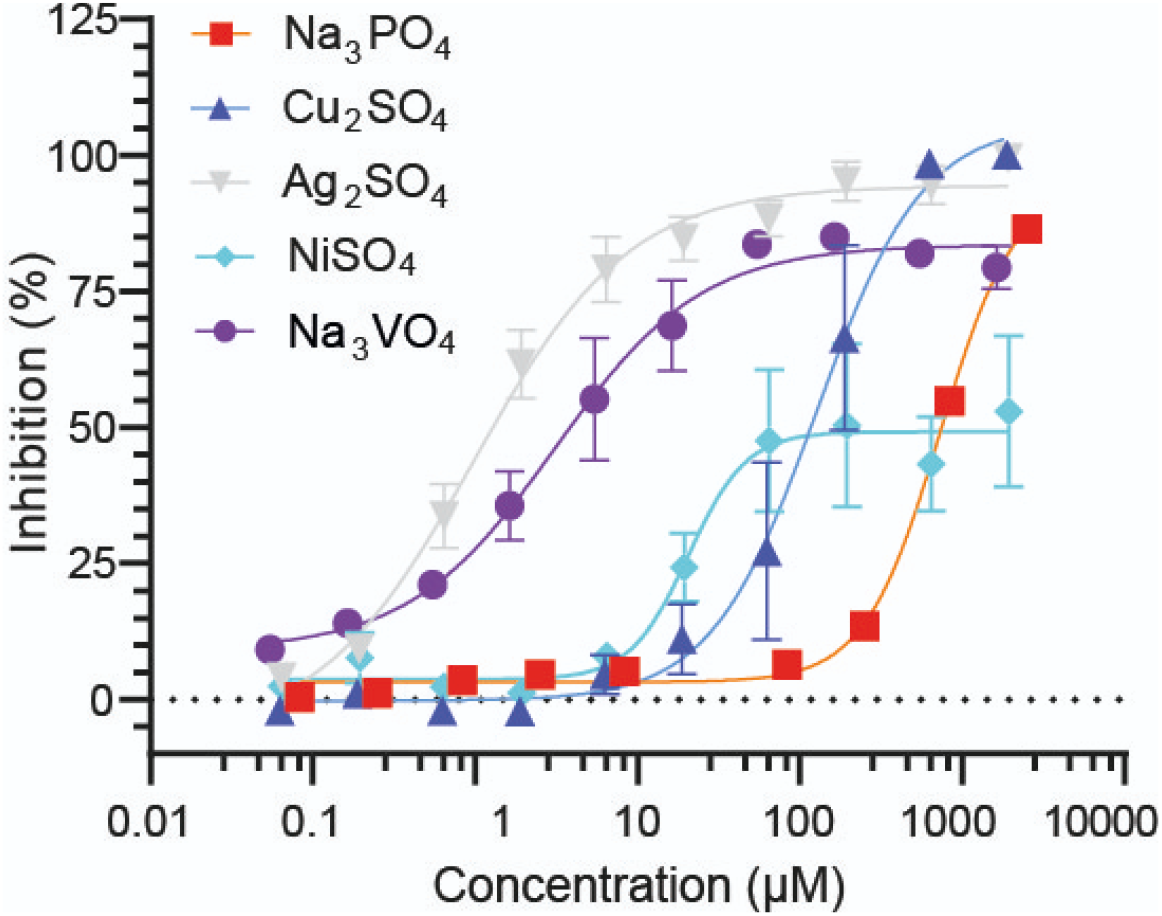
Phosphatase activity Inhibition in a concentration gradient of different salts. Concentration is expressed in μM for each compound. Fitted curves are logistic 4-parameter sigmoidal regressions.

Four salts achieved an almost complete inhibitory response (>80% of inhibition) of PP1 at concentrations superior to 200 μM. NiSO_4_ remained with an average maximum of 41%-59%. The 95% confidence interval for the reported IC50s were: [0.384 μM - 1.551 μM] for Ag_2_SO_4_, [1.597 μM - 5.373 μM] for Na_3_VO_4_, [10.01 μM - 42.41 μM] for NiSO_4_, [77.63 μM - 291.8 μM] for Cu_2_SO_4_ and [607,5 μM - 992,3 μM] for Na_3_PO_4_. Even though these chemicals were capable of interfering with the assay at low concentrations, their presence in real freshwater samples is extremely rare as discussed below.

### 3.4 Cyanobacterial lysis

#### 3.4.1 *Microcystis aeruginosa* culture

Neither cyanobacterial bloom nor MC production and release triggering conditions are totally understood (Rastogi et al., 2014). Previous works highlighted that a significant portion of the toxin produced in a bloom is retained in the intracellular compartment until cell lysis (Pham and Utsumi, 2018). Consequently, proper cellular lysis, or at least, the release of intracellular toxins, is fundamental to all methods to avoid false negatives or the underestimation of toxin concentration in the samples.

There are many lysis protocols for cyanobacteria, including complex biochemical mixtures, bead milling, and freezing, among others (Kim et al., 2009; Mehta et al., 2015). We evaluated the efficiency of a series of common or low-cost lysis protocols on a culture of *Microcystis aeruginosa*, namely: heating, freeze/thaw, sonication and filtration followed by methanol extraction. The results were compared by the MC *gold standard* quantification method, HPLC-MS. As seen in Figure 4A, when the cyanobacteria culture is fresh, the direct measuring of untreated samples (*control* treatment) reveals only a fraction of the total MC. The extracellular MC concentration (Figure 4A, ECMC), eluted in the filtration of the culture (225.03 μg/L± 82.15 μg/L), was not significantly different (T-test, *p*>0.05) to the concentration measured for the control sample without lysis treatment (156.52 μg/L ± 43.92 μg/L). All treatments performed similarly for the MC concentration detected (T-test, *p*>0.05). The determination by HPLC showed a concentration of 8225 μg/L (Supplementary Figure 3B). The methods tested, presented great accordance between them and with respect to HPLC, showing an average recovery of 92.5%, (freezing = 7291 μg/L ± 1604 μg/L, sonication = 7775 μg/L ± 846 μg/L, heat = 7763 μg/L ± 642 μg/L). However, the filtration followed by methanol extraction method described on (Sevilla et al., 2009) showed an overestimation of the total MC (TMC, sum of filtered plus extracted MC), (% recovery = 131.4%; TMC = 10809 μg/L). The intracellular, methanol-extracted concentration was 10585 μg/L ± 2027 μg/L. When we analyzed the internal organic matter release (IOMR) by absorbance 254 nm, as an index of cellular rupture, all treatments presented significantly more organic release than the control without treatment. Sonication, as expected, was the method with the highest IOMR (ANOVA, *p<0.0001*), while heating was as efficient as freezing (*p*>0.05). Notice that there is no linear increment between cell lysis and actual toxin release (Figure 4A. Right).

**Figure 4.**
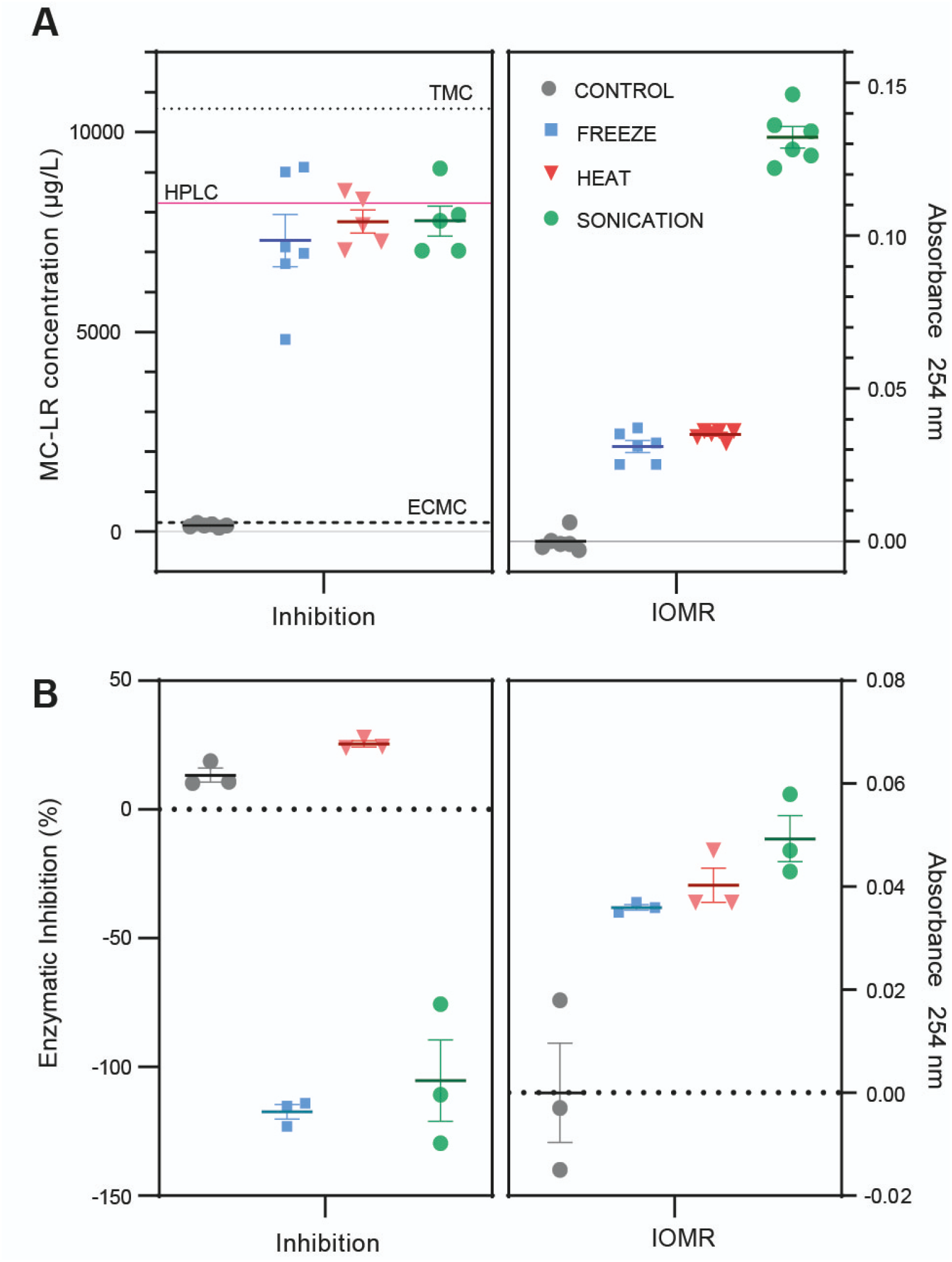
Comparison of lysis methods. A. Evaluation of lysis treatments in culture samples. *Left*: Recovered toxin concentration for each treatment. The extracellular microcystin (dashed line, ECMC = 225 μg/L) corresponds to the nylon filtered flowthrough. The total microcystin (dotted line, TMC = 10809 μg/L) corresponds to the addition of ECMC with intracellular microcystin (ICMC=10584 μg/L), obtained from the methanolic extraction of the solid algal remanent from nylon filtration with (Sevilla et al., 2009) method. The red line is the HPLC calculated concentration (8225 μg/L). *Right*: Intracellular Organic Matter Release (IOMR) for each culture, measured as UV 254 nm absorbance. IOMR control average values were subtracted from the other values. Figure 4 B. Evaluation of lysis treatments in environmental samples. *Left*: Toxin recovery evaluated as PP1 inhibition % for each treatment. *Right*: Intracellular Organic Matter Release (IOMR) for each culture, measured as UV 254 nm absorbance. IOMR values are shown relative to control values.

#### 3.4.2 Environmental Sample

We performed the same lysis protocols on a slightly toxic environmental sample, with a mid-low cyanobacteria mass (Supplementary Table 1, Sample ID#65). The IOMR and activity inhibition were measured. No dilutions were needed for the determination. The only treatment that achieved a reduction in activity (positive inhibition) due to the released MC was the heat treatment. In contrast, both freeze-thaw and sonication exhibited “negative inhibition” (Figure 4B. left). This suggests that the endogenous phosphatases of the sample exhibit a considerable effect in the inhibition and that the heating treatment inactivates these phosphatases. In contrast, the IOMR analysis showed that all the treatments release organic matter, confirming the cellular lysis. As previously seen for the culture, the highest IOMR was observed in sonication treatment, followed by heating and freeze-thawing (Figure 4B. Right). These results indicate that for complex environmental samples, the preferred lysis method is heating the sample. Furthermore, heating is reported to be a simple, economical and efficient option in comparison with high-budget lab-dependent sonication (Geada et al., 2019; Metcalf and Codd, 2000). Also, there is evidence that MC-LR tolerates exposure to high temperatures (Zhang et al., 2010). To evaluate if the concentration of MC was altered by the lysis treatment, a MC-LR calibration curve was aliquoted in 2 mL tubes and heated as previously described for lysis treatments. After that, the presence of MC-LR was measured by PP1 inhibition test. No change in MC-LR concentration with or without heating was observed when compared with the control treatment before heating (Supplementary Figure 1D).

### 3.5. Validation with environmental freshwater samples

Although PP1 is known to have sensitivity for MC, multiple reports indicate that other molecules can alter PP1 activity (Hurley et al., 2007; O’Connell et al., 2012; Peti et al., 2013). In the cellular environment, this protein interacts by more than 200 regulatory molecules. This promiscuity is detailed in the *molecular-lego* strategy and governs PP1 phosphatase target specificity (Heroes et al., 2013). Therefore, some nonspecific interaction could be present, and measurement of real water samples becomes crucial to evaluate unexpected off-target matrix effects.

#### 3.5.1 Spike recovery assays

We performed spikes of 1 μg/L, 3 μg/L and 10 μg/L in 2 mL aliquots of a low-mid density non-toxigenic algae sample. After spiking, we heated the samples according to the mentioned lysis protocol (Supplementary Table 1, Sample ID# 59). Considering the results in section *3.4.2* (Supplementary Figure 1D), no significant amount of toxin was supposed to be lost by the heat or centrifuge steps. For 10 μg/L spikes, dilutions were made. Taking into account 100% of recovery as the ideal, the spikes recovery means and SEM were calculated for each spike concentration (1 μg/L = 88.29 % ± 10.85 %, *p*=0,33; 3 μg/L = 85.78 % ± 5.974 %, *p*=0.06; 10 μg/L= 115.5 % ± 9.989%, *p*= 0.18) (Figure 5A). In all cases, no significant differences with 100% were observed for neither spike (t-test, *p*>0.05).

**Figure 5.**
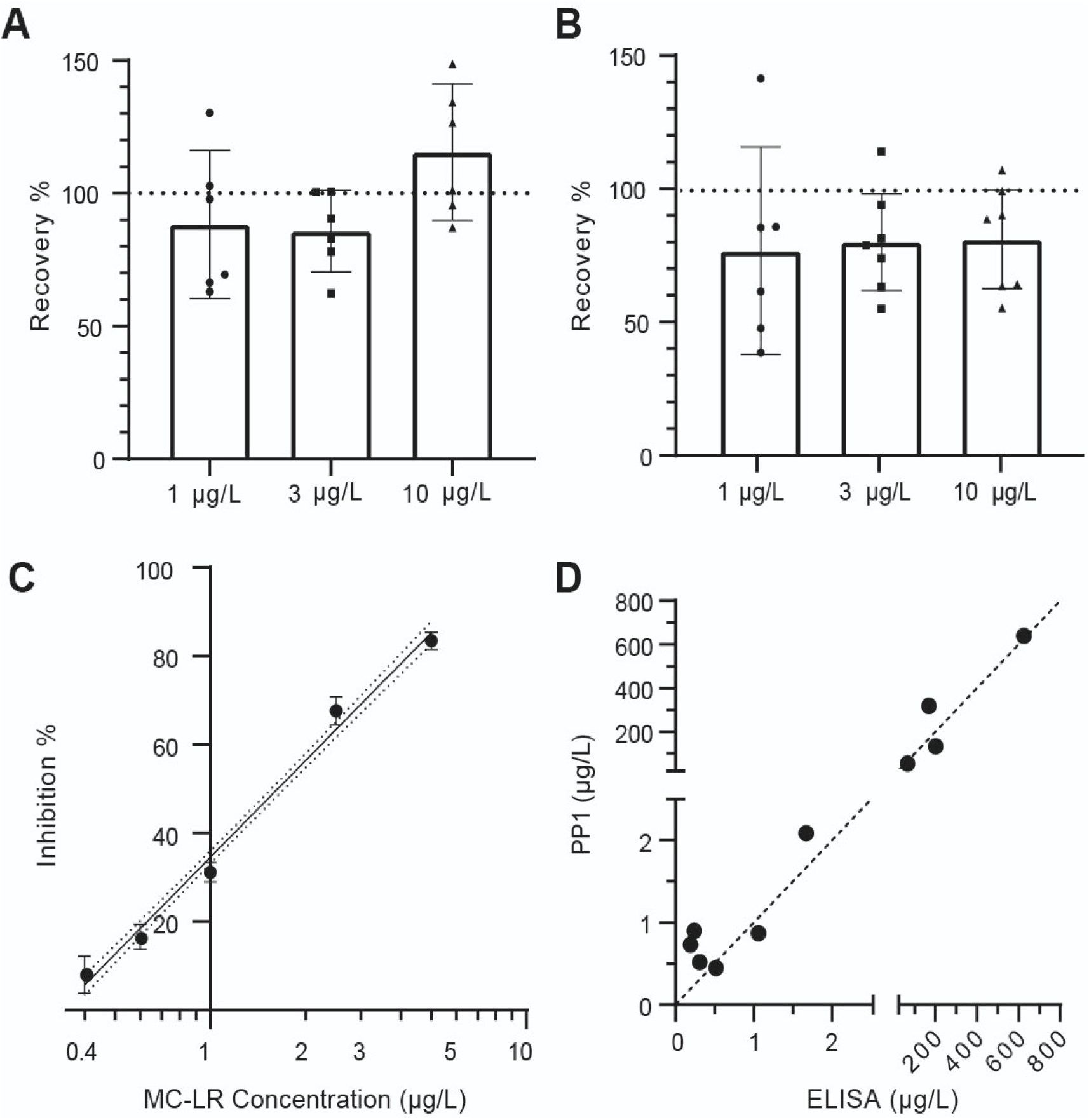
Assay Validation with environmental samples. A. Spike recovery assay of a non-toxic sample. Recovery % of the spikes of 1,3, and 10 μg MCLR/L, made to a non-toxigenic water sample (Supplementary Table 1, Sample ID# 59) (n = 6). B. Spike recovery assay of several toxic samples. Recovery % of the spikes of 1,3, and 10 μg MCLR/L, made on 7 different samples. The dotted line corresponds to the ideal 100% of recovery (in A and B). C. Inhibition calibration curve. The linear range of MCLR Inhibition of PP1 phosphatase activity is shown. Concentrations evaluated were: 0.4 ppb; 0.6 ppb; 1 ppb; 2.5 ppb; 5 ppb. The dotted line corresponds to the 95% confidence interval of the linear fit (y=72.59x + 34.51, R^2^ = 0.967). D. Correlation analysis of positive samples. Microcystin measurements are compared for both the PP1 inhibition assays and the reference ELISA values. Error bars correspond to CI95.

To study the *matrix-effect* among several different environmental samples, we also performed the spike protocol in seven other samples with a wider amount of algae density (the previously mentioned is also included in the analysis). The mean recovery and SEM for each spike concentration were calculated (1 μg/L = 77.23 % ± 15.28 %, *p*=0.19; 3 μg/L = 80.59 % ± 7.47 %, *p*=0.04; 10 μg/L = 81.62 % ± 7.62 %, *p*=0.05). Although the dispersion observed for some low concentration spikes is increased (Figure 4B), the method showed a mean recovery above 77% in all cases. Considering the variability and complexity of the water matrices, the assay was very reliable even at low MC levels.

#### 3.5.2 Validation with environmental samples

To validate the MC determination on environmental relevant samples, we received 65 freshwater samples from three different water bodies in Argentina, where active cyanobacterial monitoring programs are in progress due to frequent mild-to-severe algal blooms (Supplementary Table IDs #1-65). The samples were measured by the corresponding water authorities, with ELISA-based methods. From those 65 samples, a total of 11 officially reported to have MC concentrations above the PP1-assay detection limit in the linear range (LOD=0.4 μg/L, Figure 5C, Supplementary Table 1, Sample IDs # 12, 13, 16, 17, 18, 51, 60, 61, 62, 63, and 65). All the positive samples were also detected as positive by the PP1-assay, reporting 0 false negatives. In Figure 5D, we contrasted the MC determinations obtained by our PP1-assay vs officially reported concentrations, for the same positive water samples. The correlation between samples was calculated obtaining a Pearson correlation coefficient = 0.967 (*p*<0.0001) indicating a good correlation between measurements.

Among the 54 negative samples (reported by ELISA below 0.4 μg/L), the PP1-assay reported 6 slightly above false positives (Supplementary Table 1, Samples IDs # 34, 39, 40, 45, 46, and 47), that would imply a 0.11 false positive rate (11%) (López et al., 2015).

## Discussion

In South America and particularly in Argentina, the frequency of *CyanoHAB* events has been dramatically increasing in the last decades. Global warming, the construction of new dams, and the constant discharge of fertilizers in a region where intensive agriculture plays a predominant role in the economy, suggest that this trend will raise, with the consequent human, animal and environmental health risk (Berman, 2022; Giannuzzi and Hernando, 2022; O’Farrell et al., 2019). In Argentina, there is still no federal program for monitoring cyanobacterial blooms, possibly due to limited analytical capacity and high costs per determination (Aguilera et al., 2018)*. Microcystis* detection by microscopic cell counts or chlorophyll-A analysis is often performed by local *CyanoHABs* monitoring programs. The microscopic cell count is laborious and time-consuming and the correlation between chlorophyll-A and cyanotoxins is poor, with many cases of false positives that lead to economic losses associated with the closure of beaches and aquatic activities. Even worse are false negatives cases, with the consequent risk to the health of the exposed population (Pírez et al., 2013). Furthermore, the lack of monitoring for the presence of cyanotoxins could be underestimating the exposure of the population, as well as delaying the implementation of improvements in the water treatment and purification systems as well as remediation programs (Pírez et al., 2013; Aguilera et al., 2018). Therefore, it is of vital importance to have inexpensive and low-equipment toxin quantification methods to increase the frequency and spread of toxin detection in developing countries. Furthermore, in our region, economic instability usually causes delays in the access of imported kits, added to their high costs (Pírez et al., 2013).

In this work, we validated and optimized a PPIA. The recombinant PP1 expression and purification was improved by the coexpression with the chaperones GROEL/GROES (Peti et al., 2013). We optimized activity measurement conditions, evaluated several possible interfering substances and evaluated different storage conditions. Furthermore, we adjusted the lyophilization conditions to dry PP1 directly on the assay well. Commercially available MC kits are supplied with PP1/2A, lyophilized in a vial that requires resuspension and pipetting into the microplate. The immobilization of the protein directly in the well spares this critical and laborious step, and was previously explored by Sassolas et al (Sassolas et al., 2011) using gel entrapment.

The minimum IC50 obtained (≃ 0.3 μg/L) was consistent with the originally reported by (An and Carmichael, 1994). The mass of protein required for achieving that sensitivity in their assay (2,5-5 μg PP1/mL) is also similar to the reported herein (≃2 μg PP1/mL). However, it is possible that a lower IC50 can be achieved with other methodologies with more sensitivity, or with more active variants of PP1 (Sassolas et al., 2011). Our results suggest that part of the huge variations among IC50 (38 - 0.56 μg/L) reported in the last years (Sassolas et al., 2011; Ward et al., 1997; Wirsing et al., 1999) could be explained by differences in protein concentration or specific activity. This phenomenon was denoted by Carmichael original reports as a titration effect, also indicating that the slope of the curves increments with the increment of protein concentration (diminishing the linear range) (An and Carmichael, 1994; Carmichael and An, 1999). Heresztyn and Nicholson also performed a very rich analysis when explaining the effect of PP1 concentration in MC-LR inhibition assays, and determined a similar IC50 value with a 4-fold more sensitive technique, using phosvitin as substrate of PP1 (Heresztyn and Nicholson, 2001).

The validation performed in this work includes samples from three different water bodies and a broad range of MC levels. This is of crucial importance because of the variability associated with both, the sample itself and to sampling method and storage condition. Regarding the sample, the heterogeneity of the algae aggregates, the variable species composition, and the suspended sediment interference could lead to invalid results (Kamp et al., 2016). A limitation of freshwater evaluated in this work is the lack of samples presenting intermediate concentrations. The samples can be clustered according to concentration, one of them with very low near-background concentrations, and the other with high concentrations. Hence, the results presented here should be enriched over time with inter-institutional collaboration that will provide a wider range of MC levels in field evaluations.. The results of the 64 samples evaluated were compared with a commercial ELISA kit (Abraxis). Our PP1 method detected as positive all MC-containing samples, with no false negatives (No Type 1 Error). Furthermore, and valuable for water management authorities, the 6 false positives reported (Type 2 Error) were in most cases, slightly above the detection limit, and in all cases were below the WHO guidelines of 1 μg/L. Although the detection limits of both methods are close, the mechanism of detection is different, especially if the toxin congeners present are other than MC-LR. This could mean that some of these samples are not true “false positives”. The HPLC/MS analysis would be necessary to explore this possibility. However, even considering these 6 false positives, the specificity of 89% is an acceptable value for samples with less than 1 ug/L. These results are consistent with those reported by (Heresztyn and Nicholson, 2001), where non-toxigenic blooms showed, in some cases, a slight detection of MC.

In addition to the correct handling and homogeneity of the sample, the lysis of cyanobacteria is key to correctly measure total MC concentration. We evaluated the most common protocols to this purpose and showed that the most robust and reproducible choice is the heating method when it is associated with PPIAs. This advantage of heating was previously suggested by (Metcalf and Codd, 2000), indicating that their methodology helped in the sterilization of the biological entities of the sample. Previous authors measured field samples from reservoirs and obtained an underestimation of MC concentration. This phenomenon can be explained by their lysis methodology: they freeze-thawed the samples and measured them by PP1 assay (Heresztyn and Nicholson, 2001). We hypothesized that endogenous phosphatase activity of an environmental sample can heavily distort the results obtained, as shown in our lysis analysis performed with an environmental sample (Figure 4B). If this is the case, sonicated and freeze/thawed showed “negative inhibition” (more phosphatase activity) than the control treatment because of the phosphatase activity present in the samples. Moreover, the obtained IOMR profile both in the *Microcistys* culture and environmental samples indicate that the freezing and sonication are lysing the cyanobacteria. In the case of the culture, the MC concentration was extremely high (Figure 4A), meaning that to quantify it correctly, we needed to make dilutions up to 1/5000-fold. This dilution also diminished the *matrix effect*, including the endogenous active phosphatases in the sample. When working with environmental samples, this fact becomes more relevant, because the amount of MC-LR production is usually significantly lower, and so are the dilutions needed, leading to more endogenous active phosphatases. Other methods such as ELISA can also be affected by cellular matrix but would not be influenced by the release of endogenous phosphatases making the lysis by freeze-thaw suitable for this methodology. To conclude, heating seems like the most economical and viable option that, in addition to releasing the toxins, inactivates endogenous phosphatases.

The matrices of environmental samples can be extremely variable both in chemical composition and conductivity. The assay showed a very good robustness in the freshwater ranges of salt and pH. Also, the construction of inhibition curves for a diverse spectrum of anions and cations allows us to assume that the salts that are usually present in fresh water will not interfere with the assay, at least in the expected reported levels. Of special interest is the case of coordination metals like Mg^+2^, that did not present any inhibition in the evaluated range. Also, the reaction was not significantly altered by phosphate salts up to 100 μM (more than the average natural of 0.05-5 μM) (Hudson et al., 2000). However, four of them showed a significant inhibition in low levels, namely Cu ^+2^, Ag^+1^, Ni^+2^ and VO_4_^−3^ and worth a brief discussion. Both Ag^+1^ and Cu^+2^, are used in a patented chemical-based algal lysis protocol (RAPID ANALYSIS FOR CYANOBACTERIAL TOXINS, US Patent 9,739,777 B1). Our results indicate that this method would not be suitable for PPIAs. Silver nanoparticles can catalyze the reduction reaction of nitrophenol (yellow) to aminophenol (colorless) (Ai and Jiang, 2013) in a reducing medium like the one used in PPIA. This could explain the extremely sensitive silver inhibition, involving only the product, but without Ag-protein interaction. Concerning copper, there is evidence that can be coordinated by PP1*γ* structure in a non-active form (Miyazaki et al., 2009). However, it exhibited inhibition at concentrations higher than present in nature ≃ 0.003-0.38 μg/L (Australian and New Zealand Environment and Conservation Council - ANZECC, 2000, pt. 8.3.2). The cations Ag^+1^, Ni^+2^, are rarely present in freshwater (less than 1 μg/L). In the case of VO_4_^−3^, we had previously tested groundwater samples and detected basal inhibition (approximately 20%). Correlation analysis showed a slight relation with arsenic and fluor concentration (Pearson = 0.79, *corrplot R package*). Interestingly, arsenic and fluor salts have shown no significant effect on the inhibition curves (Supplementary Figure 2). However, F and As usually are geologically related to different forms of V (Puntoriero et al., 2015), being the most prevalent form VO_4_^−3^ (Gustafsson, 2019). It is known that VO_4_^−3^ can inhibit other types of phosphatases, the PTPs (Protein Tyrosine Phosphatases). In the ’80s, (Swarup et al., 1982) showed that VO_4_^−3^ selectively inhibits the Protein Tyrosine Phosphatases but not Serine-Threonine Protein Phosphatases -as PP1-activity with radiolabeled histones,. Another work (MacKintosh et al., 1996) showed that some *E. coli*-produced PP1*γ* presented some amount of Tyrosine phosphatase activity that could be inhibited by VO_4_^−3^. In accordance with MacKintosh, our *E.coli*-produced PP1α presented significant VO_4_^−3^ inhibition in the pNPP phosphatase activity. Beyond the biochemical relevance for understanding PP1 chemistry, the presence of VO4^−3^ in freshwater is rare.

## Conclusion

In this work we optimize a microplate-based assay for the determination of MC mediated by the inhibition of recombinantly expressed PP1. The assay proved to be robust, reproducible and sensitive (LOD 0.4 ug/L), allowing the detection of MC below the WHO guideline of 1ug/L. Lysis by heating the sample proved to be the method of choice for the quantification of MC by PPIAs. The assay was validated with samples from dams of Argentina, showing a high degree of correlation with the results obtained by a commercial ELISA kit. The sample method described here is low-cost and requires minimal equipment, making it suitable to expand the monitoring capabilities of developing countries.

## Conflict of interest

Authors declare no conflict of interest

## Acknowledgments

We want to acknowledge all the people that directly or indirectly collaborated with the development of this work. The chaperone plasmid was kindly provided by Dr Diana Wetzler from Universidad de Buenos Aires. Also, acknowledge Domingo Tagliafico from Soluciones Analíticas S.A, for his collaboration in the use of Thermo Scientific™ Vanquish™ HPLC system. The environmental samples were gently provided by our collaborators in water management organizations. Among them; Facundo Bordet from Comisión Técnica Mixta de Salto Grande, Marcia Ruiz from Instituto Nacional del Agua - CIRSA, Ayelen Othaz from Autoridad Interjurisdiccional de Cuenca and Ariana Rossen from Instituto Nacional del Agua, Monica Cavallo from Autoridad del Agua de la Provincia de Buenos Aires. Finally we want to thank for the received support and valuable opinions of our colleagues Yamila Gandola and Macarena Alvarez, Javier Santos, Ignacio Castro from Universidad de Buenos Aires, Fernando Macario from Fundación INVAP and Leonardo Sierra from IHLLA.

## Abbreviations

PP1: protein phosphatase 1
MC: microcystin
MC-LR: microcystin LR
*CyanoHABs*: cyanobacterial harmful algal blooms
IOMR: internal organic matter release
DTT: ditiotreitol
ELISA: enzyme-linked immunosorbent assay
TMC: Total microcystin
ECMC: Extracellular microcystin
ICMC: Intracellular microcystin
HESI: Heated electrospray ionization
SEM: Standard error of the mean

## Notes

### Competing Interest Statement

The authors have declared no competing interest.

