## Supplementary for "Optimization and validation of a Protein Phosphatase inhibition assay for accessible microcystin detection"

***Supplementary material:***

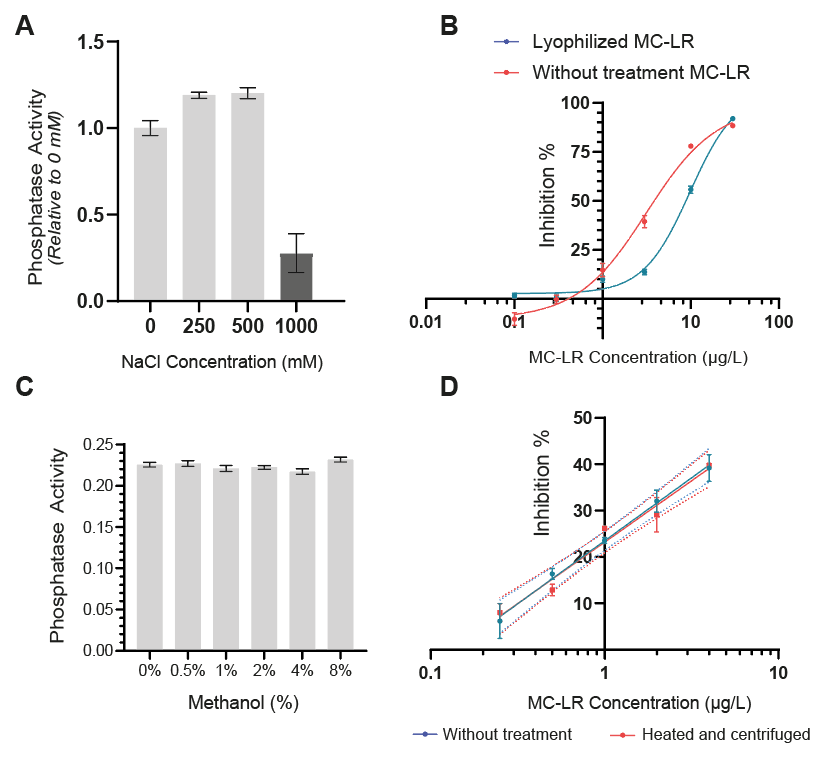

**Sup. Fig. 1. Stability of PPI response and MC inhibition under different conditions.** A. High salinity phosphatase inhibition: Phosphatase activity was measured in the presence of a gradient of NaCl (250-1000 mM). In concentrations higher than 1 M the activity significantly diminished. (n=2).

B. Lyophilization of protein with MC-LR: The protocol of lyophilization was tried by adding both protein and MC before lyophilization. When inhibition was tested in comparison with the non-lyophilized toxin, the resulting inhibition was significantly lower (p<0.05). C. Methanol does not affect phosphatase inhibition: Phosphatase activity in the presence of 100 μl of a gradient of the methanol-water mixture. There is no observable inhibition of activity up to 8% of methanol. D. Microcystin is not being degraded by heating: PP1 microcystin-LR inhibition curve, before and after being heated 15 minutes at 95°C and centrifuged to maximum speed for 15 minutes. The resulting inhibition indicates no significant difference either in slope or ordinate of the curve fit (N=2. p>0.05). This suggests that there is no degradation or precipitation of MC due to heat or centrifugation. Error bars correspond to SD. Dotted lines correspond to the 95% confidence interval of the linear fitting.

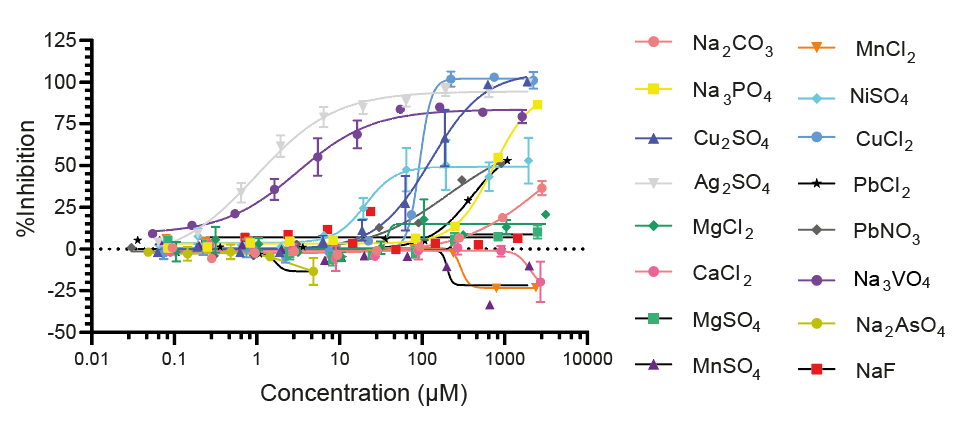

**Sup. Fig. 2:** Activity Inhibition curves of all 16 salts tested (most of them) in the range from 10 μg/L to 300 mg/L (of the mentioned metal). Each curve is an average of *n* independent gradient inhibition assays. Furthermore, each assay was performed with 3 technical repeats for each concentration. (n=3).

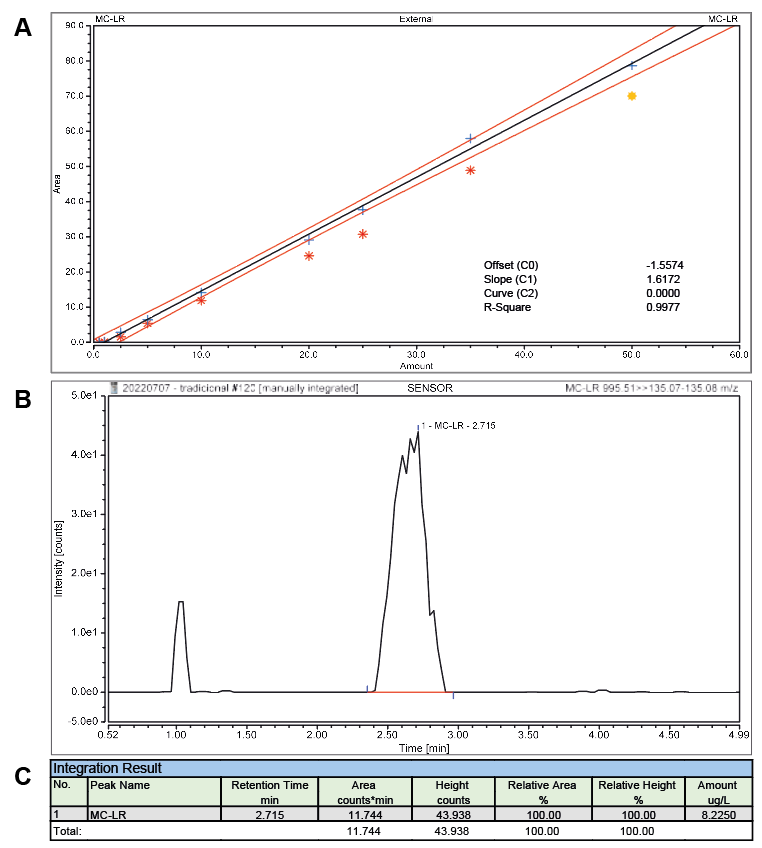

**Sup. Fig. 3. Quantification of MC-LR by gold standard method HPLC-MS.** A. MC-LR calibration curve. A MC-LR calibration curve was carried out using five levels, Abraxis™ MC-LR Standard solution (0.5, 1.0, 2.5, 5.0, 10.0, 20.0, 25.0, 35.0, 50.0µg/L); R-Square: 0.9977. B. MC-LR peak. The MC-LR identification was performed by comparing the acquired mass spectrum and retention time to the reference spectrum and retention time for calibration standard (MCLR 995.51>>135.07-135.08 m/z). C. Integration Result. The Table shows important quantification parameters as retention time (min), area (counts*min) and amount (µg/L) by Software in use Chromeleon 7.3.0.60919
**Sup. Table 1:** Water samples used in this work for validation, spikes and lysis protocol.

| **ID** | **Sampling Date** | **Origin** | **PP1 (μg/L)** | **ELISA (μg/L)** |
| --- | --- | --- | --- | --- |
| 1 | 7/7/21 | El Chocon dam, Neuquen province, Argentina | ND | ND |
| 2 | 7/7/21 | El Chocon dam, Neuquen province, Argentina | ND | ND |
| 3 | 7/7/21 | El Chocon dam, Neuquen province, Argentina | ND | ND |
| 4 | 7/7/21 | El Chocon dam, Neuquen province, Argentina | ND | ND |
| 5 | 7/7/21 | El Chocon dam, Neuquen province, Argentina | ND | ND |
| 6 | 13/07/2021 | Salto Grande dam, Entre Rios province, Argentina | ND | ND |
| 7 | 13/07/2021 | Salto Grande dam, Entre Rios province, Argentina | ND | ND |
| 8 | 21/4/22 | Buenos Aires province, Argentina | ND | 0.16 |
| 9 | 21/4/22 | Buenos Aires province, Argentina | ND | <0.15 |
| 10 | 21/4/22 | Buenos Aires province, Argentina | ND | <0.15 |
| 11 | 21/4/22 | Buenos Aires province, Argentina | ND | <0.15 |
| 12 | 21/4/22 | Buenos Aires province, Argentina | 0.87 | 1.06 |
| 13 | 21/4/22 | Buenos Aires province, Argentina | 0.45 | 0.52 |
| 14 | 21/4/22 | Buenos Aires province, Argentina | 0.29 | <0.15 |
| 15 | 21/4/22 | Buenos Aires province, Argentina | 0.39 | 0.15 |
| 16 | 21/4/22 | Buenos Aires province, Argentina | 0.52 | 0.31 |
| 17 | 21/4/22 | Buenos Aires province, Argentina | 0.73 | 0.19 |
| 18 | 21/4/22 | Buenos Aires province, Argentina | 0.90 | 0.24 |
| 19 | 9/5/22 | Río de la Plata, Buenos Aires province, Argentina | ND | ND |
| 20 | 9/5/22 | Río de la Plata, Buenos Aires province, Argentina | ND | ND |
| 21 | 9/5/22 | Río de la Plata, Buenos Aires province, Argentina | ND | ND |
| 22 | 9/5/22 | Río de la Plata, Buenos Aires province, Argentina | ND | ND |
| 23 | 9/5/22 | Río de la Plata, Buenos Aires province, Argentina | ND | ND |
| 24 | 9/5/22 | Río de la Plata, Buenos Aires province, Argentina | ND | ND |
| 25 | 9/5/22 | Río de la Plata, Buenos Aires province, Argentina | ND | ND |
| 26 | 9/5/22 | Río de la Plata, Buenos Aires province, Argentina | ND | ND |
| 27 | 9/5/22 | Río de la Plata, Buenos Aires province, Argentina | ND | ND |
| 28 | 9/5/22 | Río de la Plata, Buenos Aires province, Argentina | ND | ND |
| 29 | 9/5/22 | Río de la Plata, Buenos Aires province, Argentina | ND | ND |
| 30 | 9/5/22 | Río de la Plata, Buenos Aires province, Argentina | ND | ND |
| 31 | 9/5/22 | Río de la Plata, Buenos Aires province, Argentina | ND | ND |
| 32 | 9/5/22 | Río de la Plata, Buenos Aires province, Argentina | ND | ND |
| 33 | 9/5/22 | Río de la Plata, Buenos Aires province, Argentina | ND | ND |
| 34 | 9/5/22 | Río de la Plata, Buenos Aires province, Argentina | 0.73 | ND |
| 35 | 9/5/22 | Río de la Plata, Buenos Aires province, Argentina | 0.27 | ND |
| 36 | 9/5/22 | Río de la Plata, Buenos Aires province, Argentina | 0.33 | ND |
| 37 | 9/5/22 | Río de la Plata, Buenos Aires province, Argentina | ND | ND |
| 38 | 9/5/22 | Río de la Plata, Buenos Aires province, Argentina | 0.37 | ND |
| 39 | 9/5/22 | Río de la Plata, Buenos Aires province, Argentina | 0.44 | ND |
| 40 | 9/5/22 | Río de la Plata, Buenos Aires province, Argentina | 0.48 | ND |
| 41 | 9/5/22 | Río de la Plata, Buenos Aires province, Argentina | 0.30 | ND |
| 42 | 9/5/22 | Río de la Plata, Buenos Aires province, Argentina | ND | ND |
| 43 | 9/5/22 | Río de la Plata, Buenos Aires province, Argentina | 0.40 | ND |
| 44 | 9/5/22 | Río de la Plata, Buenos Aires province, Argentina | 0.38 | ND |
| 45 | 9/5/22 | Río de la Plata, Buenos Aires province, Argentina | 0.43 | ND |
| 46 | 9/5/22 | Río de la Plata, Buenos Aires province, Argentina | 0.74 | ND |
| 47 | 9/5/22 | Río de la Plata, Buenos Aires province, Argentina | 0.41 | ND |
| 48 | 9/5/22 | Río de la Plata, Buenos Aires province, Argentina | 0.28 | ND |
| 49 | 9/5/22 | Río de la Plata, Buenos Aires province, Argentina | ND | ND |
| 50 | 9/5/22 | Río de la Plata, Buenos Aires province, Argentina | ND | ND |
| 51 | 9/5/22 | Río de la Plata, Buenos Aires province, Argentina | 2.09 | 1.67 |
| 52 | 9/5/22 | Río de la Plata, Buenos Aires province, Argentina | ND | ND |
| 53 | 9/5/22 | Río de la Plata, Buenos Aires province, Argentina | ND | ND |
| 54 | 9/5/22 | Río de la Plata, Buenos Aires province, Argentina | ND | ND |
| 55 | 9/5/22 | Río de la Plata, Buenos Aires province, Argentina | 0.25 | ND |
| 56 | 9/5/22 | Río de la Plata, Buenos Aires province, Argentina | ND | ND |
| 57 | 9/5/22 | Río de la Plata, Buenos Aires province, Argentina | ND | ND |
| 58 | 9/5/22 | Río de la Plata, Buenos Aires province, Argentina | ND | ND |
| 59 | 2/6/22 | Salto Grande dam, Entre Rios province, Argentina | 0.26 | <0.15 |
| 60 | 2/6/22 | Salto Grande dam, Entre Rios province, Argentina | 134.65 | 201 ppb |
| 61 | 2/6/22 | Salto Grande dam, Entre Rios province, Argentina | 319.10 | 170 ppb |
| 62 | 2/6/22 | Salto Grande dam, Entre Rios province, Argentina | 638.71 | 625 |
| 63 | 2/6/22 | Salto Grande dam, Entre Rios province, Argentina | 54.70 | 65.5 |
| 64 | 2/6/22 | Salto Grande dam, Entre Rios province, Argentina | 0.29 | <0.15 |
| 65 | 4/05/22 | Pond in Buenos Aires province, Argentina | 0.90 | 0.61 |
